# Effect of alcohol on the speed of shifting endogenous and exogenous attention

**DOI:** 10.1101/2024.04.24.590923

**Authors:** Alexander Thiele, Christopher Heath, Jessica Kate Pearson, Sidharth Sanjeev, Jenny C. A. Read

## Abstract

The study aimed to investigate to what extent acute moderate doses of alcohol affect the speed of endogenous versus exogenous attentional shift times. Subjects viewed an array of 10 moving clocks and reported the time a clock indicated when cued. Target clocks were indicated by peripheral or central cues, including conditions of pre-cuing. This allowed assessing shift times when attention was pre-allocated, when peripheral cues triggered exogenous attention shifts, and when central cues triggered endogenous attention shifts. Each subject participated in 2 sessions (alcohol/placebo), whereby the order of drug intake was counterbalanced across subjects, and subjects were blinded to conditions. Confirming previous results, we show that pre-cuing resulted in the fastest shift times, followed by exogenous cuing, with endogenous attentional shifts being slowest. Alcohol increased attentional shift times across all 3 conditions compared to placebo. Thus, the detrimental effects of alcohol on attentional shift times did not depend on the type of attention probed.

## Introduction

Multiple studies have explored how quickly covert attention can be shifted between different locations, dating back to experiments performed by Wilhelm Wundt using a complication clock apparatus (Wundt, 1883). Since then, a variety of techniques have been employed to determine the speed of attentional shifts, including orienting paradigms (Carlson, Hogendoorn, & Verstraten, 2006; M. I. Posner, 1980; Yantis & Jonides, 1990), paradigms investigating attentional gating (Reeves & Sperling, 1986) and visual search paradigms (Wolfe, Alvarez, & Horowitz, 2000). An emerging pattern from all those studies is that two types of attention can be dissociated, namely bottom-up and top-down attention (Carlson et al., 2006; Duncan, 1984; M. I. Posner, 1980; M.I. Posner & Petersen, 1990; Theeuwes, 1991). While bottom-up attention is reflexive and triggered by salient unexpected stimuli in the external world (often peripheral cues in experimental situations), top-down attention is wilful, and can be triggered by central cues (which often need to be interpreted in experimental situations). In addition to the different mechanisms by which bottom-up and top-down attention is triggered, they also differ in terms of how quickly they can be allocated (shifted) to the relevant stimuli. Bottom-up attention shifts are faster than top-down attention shifts by ∼50-100ms (Carlson et al., 2006; Chakravarthi & VanRullen, 2011; Muller & Rabbitt, 1989; M. I. Posner, 1980).

Alcohol affects several aspects of cognition, including divided and covert attention (Schulte, Muller-Oehring, Strasburger, Warzel, & Sabel, 2001). At low dosage (0.33ml/kg body weight) it can improve accuracy on four-choice serial reaction time and on visual search tasks, while at high dosage (1.0ml/kg body weight) it increases reaction times and decreases accuracy (Maylor & Rabbitt, 1987). Moreover, it affects speed accuracy trade-offs in 4-choice reaction time, number pairs and visual search tasks, as well as negatively affecting working memory (Benson, Tiplady, & Scholey, 2019). Moderate levels of alcohol reduce the speed of temporal visual processing by increasing neuronal latencies across a variety of visual tasks (Khan & Timney, 2007). It has been argued that significant perceptual, motor and cognitive impairments occur even at very low blood alcohol levels (<0.02gm/100ml) (Koelega, 1995; Ogden & Moskowitz, 2004).

One of the mechanisms by which alcohol might negatively affect attention, is through reduction of inhibitory control. Where cognitive resources need to be directed towards relevant stimuli, the inhibition of irrelevant stimulus interference is an important part of attentional selection (Fox, 1995; Klein, 2000). Abroms et al. (Abroms, Gottlob, & Fillmore, 2006) found that intentional inhibitory control over selective attention was negatively affected by alcohol intake while automatic mechanisms were not. This suggest that top-down (voluntary) attention shift times might be increased by alcohol, while bottom-up (automatic) attention shift times might be unaffected or at least be less affected by moderate doses of alcohol. A similar prediction would also be made based on the global-slowing hypothesis (Maylor & Rabbitt, 1993). It predicts that alcohol induced increases in attention shift times are proportional to the speed of the component process under control conditions, with a proportionality constant >1. Since the baseline shift times for bottom-up attention are faster, the absolute alcohol induced increase would be smaller for bottom-up attention. However, Ryan et al. (Ryan, Russo, & Greeley, 1996), did not find evidence for the global-slowing hypothesis. Finally, it has been suggested that the effect of alcohol increases with task complexity (Maylor & Rabbitt, 1993), whereby exogenously cued attention shifts (a low level automatic task) have been reported to be unaffected by moderate levels of attention (Post, Chaderjian, & Maddock, 2000).

To test the prediction that endogenous attention shifts are more strongly affected by moderate levels of alcohol, we used a method of estimation of attention shift speed, based on a complication-clock apparatus originally used by Wilhelm Wundt (Wundt in Carlson et al., 2006). It involves participants watching a fixation point surrounded by moving clocks. After a clock is cued, participants shift their attention to that clock and record the position of the hand they perceived on the clock when the clock was cued.

## Methods

### Participants

Twenty participants took part in the experiment. Participants were undergraduate volunteers from Newcastle University. The participants included seven males and thirteen females, aged from 19 to 22. All had normal or corrected-to-normal vision and no previous alcohol problems. Participants were instructed not to have consumed alcohol 12 hours prior to the experiment. Equally, they were instructed that they need to have abstained from recreational drugs use for at least 24 hours before they participate in a session.

Subjects had to perform the task twice, whereby each session had to be performed on separate days (at about the same time of a day, i.e. if a subject did session one in the morning, they performed session 2 on a different day, but also in the morning. They were also informed prior to the session that alcohol may be consumed during one of the sessions, whereby they were blinded to which this was. They were therefore instructed not to drive or operate any motorised equipment for up to 6 hours after a session and to ensure they had no plans to return to lectures or work after participation. All participants gave informed consent prior to participating in the experiment. Ethical approval was obtained from the ethics committee of the Faculty of Medical Sciences, Newcastle University (00860/2015).

### Apparatus/ Materials

The experiment was run on a Windows PC running MATLAB 2015a (62-bit) (The MathWorks, Inc.) using PsychToolbox extentions (Brainard, 1997). The stimuli were displayed on an Iiyama 22” monitor (100 Hz, 2048 x 1536 resolution) that was controlled by the PC. Observers sat with their chin on a chin rest 60cm away from the monitor. Responses were collected from the keyboard, using the left and right arrow keys to move the clock hand and the space bar to confirm the participant’s observed time. Prior to the experiment, participants were provided with an information sheet explaining the psychophysical task. They were required to sign a consent form to participate in the experiment. The consent form gave information about the task, the detailed procedures, the purpose of the study, how data would be used and anonymized. Here they also confirmed that they had not eaten for 2 hours before the experiment or drunk alcohol for 12 hours before the experiment. The form contained a second section where participants had to sign (after completion of an experimental session) to confirm that they were able to leave the lab and carry on with their daily routines independently.

#### Application of alcohol

Subjects were either given an alcoholic or a non-alcoholic beverage before each session (whereby the order of drug/no drug was counterbalanced across subjects). The alcoholic cocktail used in the study was a modified version of the Brazilian cocktail named Caipirinha. The ingredients included Cachaça (40% alcohol content), lime, sugar and orange juice. The UK legal limit for drivers is 80 mg of alcohol per 100 ml of blood, commonly referred to as BAC (blood alcohol concentration). In terms of breath alcohol this is 350 μg (micrograms) per litre (which is now the usual official measure in the UK). To calculate the amount of Cachaça to include in each Caipirinha cocktail to bring the participants BAC to 80 mg of alcohol per 100 ml of blood, the Widmark formula was used:

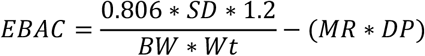

The Widmark formula estimates peak blood alcohol concentration (EBAC), a variation, including drinking period in hours. Where:

- 0.806 is a constant for body water in the blood (mean 80.6%),
- SD is the number of standard drinks containing 17 grams of ethanol,
- 1.2 is a factor to convert the amount in grams to Swedish standards set by The Swedish National Institute of Public Health,
- BW is a body water constant (0.58 for men and 0.49 for women),
- Wt is body weight (kilogram),
- MR is the metabolism constant (female = 0.017, male = 0.015) and
- DP is the drinking period in hours.

Using the criteria two different formulas for each gender were derived to give the amount (ml) of 100% alcohol needed to get the participant to 0.08% alcohol blood level:

- Male X = (17*(0.08+0.017)*0.58*Bodyweight (kg))/(0.806*1.2)
- Female X= (17*(0.08+0.017)*0.49*Bodyweight (kg))/(0.806*1.2)

Cachaça has an alcohol content of 40%, therefore X was divided by 0.4, to give the amount of Cachaça needed for the alcoholic drink. The non-alcoholic placebo cocktail used the same overall drink volume (200ml) but added rum aroma instead of Cachaça (1 volume of rum aroma to 2 volumes of rum), to mimic the taste.

An Alcohawk Pro Digital Breathalyser Alcohol Detector was used to measure breath alcohol content (BrAC, details below). The breathalyser recordings were used as an indirect measure of participants’ blood alcohol levels.

### Design

The design of the experiment was within-subject repeated measurements. Each participant carried out the experimental and placebo conditions in separate sessions. Participants were blinded to the condition. The two sessions were performed on different days, but at roughly the same time of day (the second session start time was always within two hours of the start time of the first session-for example, if the first session took place at 11am, the second session would take place between 9am and 1pm on a subsequent day) to control for effects of circadian rhythm on attention. Approximately half of participants performed the experimental condition first and the other half performed the placebo condition first.

### Procedure

Before the experiment, participants were given the experiment information sheet and questionnaire via email. They had to fill out the pre-experiment questionnaire and email it to the experimenter. Upon arrival for the first session, they were given a consent form to read and sign. They were then weighed (to allow the amount of alcohol they would need to consume to be calculated) and breathalysed (to ensure they had no alcohol in their systems). They were briefed again about the experiment, i.e. the psychophysical task. Any questions about the task were then answered. This was followed by a 90-trial practice session, where they got used to the task, and obtained initial training. The participant then drank the cocktail prepared by the experimenter. A 25 minute waiting period occurred after the participant had finished the cocktail to allow the alcohol to be metabolised and take effect. Participants had the option of waiting in the experiment room or leaving and returning at the end of the waiting period. The participant was then breathalysed again, before performing the experimental set of 150 trials. The participant was breathalysed again post-experiment. In the second session, the same procedure was followed, but the participant was not weighed and did not perform the practice set of trials.

The psychophysical task was as described in (Carlson et al., 2006). It consisted of 3 different attention conditions (pre-cue, exogenous and endogenous cuing), where subjects had to read the time on one of 10 possible clocks, when the relevant clock was cued (see below). Each condition occurred 50 times, randomly selected without replacement. Each trial started with a central fixation spot, which subjects were asked to fixate. Before the start of the trial the computer randomly assigned which clock would be cued on that given trial. In the pre-cue conditions the fixation spot appeared along with a central cue, a short line (length: 2 deg of visual angle (dva)), emanating from close to the fixation cross, pointing to the future relevant clock location. After 1 second, 10 clock faces (∼2.5 dva diameter) appeared simultaneously, arranged in a circle at an eccentricity of 7 dva on the screen (see Figure 1). Each clock showed a single clock hand with a randomly allocated starting position. The clock hands rotated at 1Hz, i.e. one revolution per second. At a randomly selected time after the start of the trial (drawn from a uniform distribution from 100-1000ms after clock onset, steps of 10ms) one of the clocks would be cued (details below). Shortly after the cue (500ms), the clocks were replaced by one central clock. Participants were required to indicate the position of the hand on the cued clock at the time of the cue by moving the hand of the central clock to the same position using the left/right arrow buttons of the keyboard. There were 3 different conditions in the task; in the exogenous cue condition, the rim of the cued clock changed from black to red at the time of cuing for 5 frames (50ms). In the endogenous cue condition, a line appeared emanating from the fixation point pointing towards one of the clocks (for 5 frames, i.e. 50 ms). In the baseline (pre-cue) condition, a line appeared before the clock faces (as described above), indicating the position of the clock that would be cued. That specific clock was then cued during the trial as in the exogenous cuing condition. Figure 1A, B, and C illustrate the progression of events in each of the 3 conditions (see also the videos in Carlson et al., 2006).

**Figure 1:**
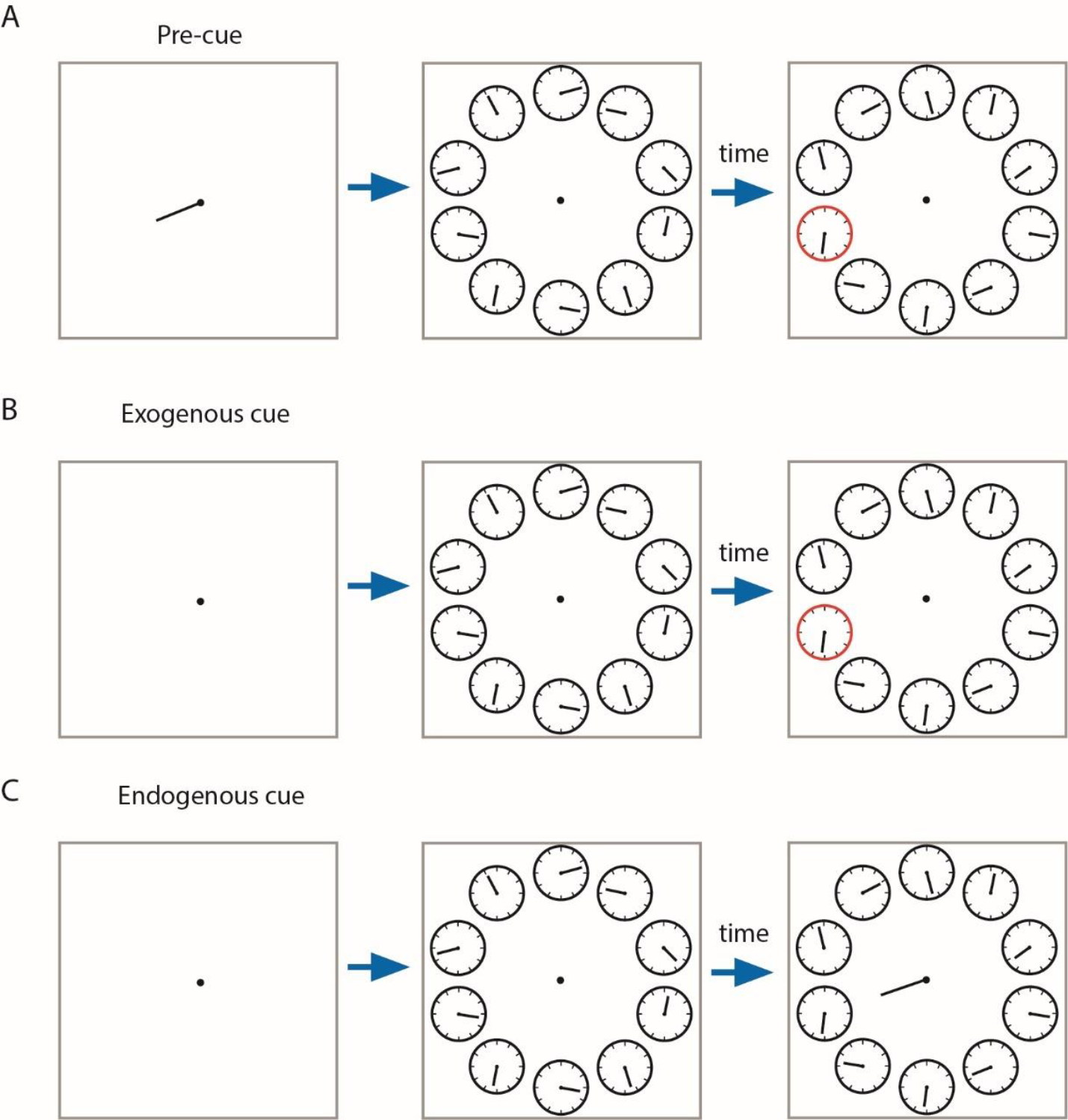
Experimental setup for the 3 attention conditions. A) The top row indicates the events that occur in the pre-cue condition, B) middle row the events in the exogenous condition, and C) bottom row the events that occur in the endogenous cuing condition. The blue arrow indicates time passing. This was fixed for the first period, but variable for the second period.

### Data Analysis

The latency between the time of cueing and the time of cueing reported by the participant (i.e. the time indicated by the adjusted the clock hand) was calculated for each trial. This could result in latencies of 0-999ms (one full revolution of the clock hand) in principle. However, latencies that exceeded 900 ms were calculated as latency-1000. This cutoff latency was accepted as it allows for some random perceptual/memory errors to be distributed around the actual time that was present at cuing. This resulted in all final latencies to range from -99ms to 900ms. The mean latency for the 3 cuing and the 2 experimental (drug/no drug) conditions were then calculated for each participant. A mixed model ANOVA based on the subject means determined whether cuing condition (3 levels, within subject repetition), experimental condition (2 levels, within subject repetition) or the order of the experimental condition (alcohol or placebo in first session, between subject factor) significantly affected the attentional shift times.

## Results

Twenty participants performed the study, whereby 9 participants took the non-alcoholic drink first, and the remaining eleven took the alcoholic drink before the first session (order of experimental conditions). Confirming previous results (Carlson et al., 2006), we found that shift times were fastest in the pre-cue condition, followed by the exogenous cuing, and slowest in the endogenous cuing condition (Figure 2 and Table 1).

**Table 1.** Mean attentional shifting times in milliseconds (ms) for the placebo and experimental conditions for each of the 3 cueing conditions. Mean shift times were first calculated for each subject. Reported here are group means and SEM obtained from subject means.

| Condition | Mean attentional shifting times (ms) | Standard Error | Difference (ms) |
| --- | --- | --- | --- |
| Pre, Placebo | 74.23 | 9.00 | 45.53 |
| Pre, Alcohol | 119.76 | 16.19 |  |
| Exogenous, Placebo | 191.77 | 13.96 | 32.34 |
| Exogenous, Alcohol | 224.11 | 15.96 |  |
| Endogenous, Placebo | 314.94 | 15.52 | 29.46 |
| Endogenous, Alcohol | 344.40 | 15.40 |  |

**Figure 2:**
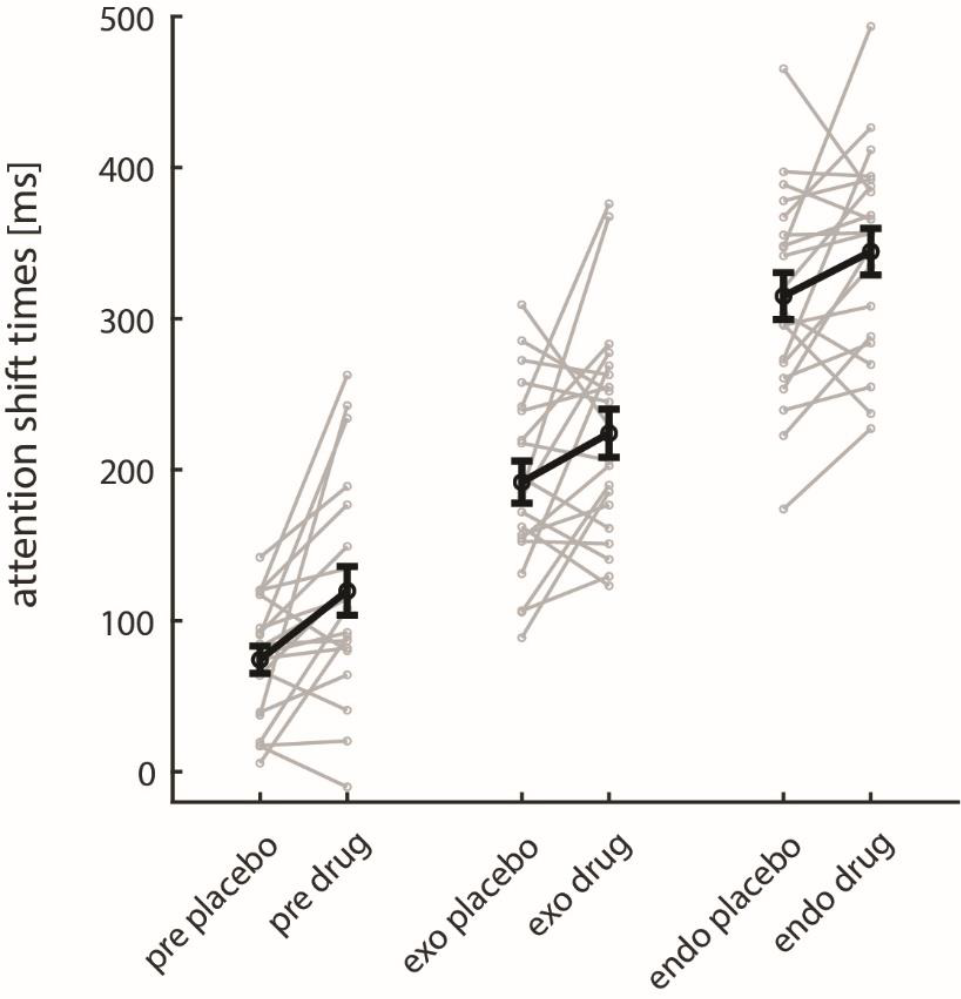
Attentional shift times for the 3 attention conditions when subjects were not under the influence of alcohol (placebo) and when they were under the influence of alcohol (drug). The three cuing conditions are separated along the x-axis, grey data points and lines give data for individual subjects, black data points show mean attentional shift times along with S.E.M.

In Figure 2 it can be seen that mean attention shift times were higher for all cueing conditions when alcohol (drug) was consumed, but the individual data points (grey symbols and linking lines) show that some subjects were faster under alcohol compared to non-alcohol (placebo) conditions. This can be explained by the order in which drug/placebo sessions were arranged and the effect of learning between session, which will be explained in more detail later. Table 1 shows the values of the mean shift times, along with differences between alcohol and placebo conditions. Difference values were larger for pre-cue conditions than other conditions, but condition dependent difference were not significant as revealed by a mixed model ANOVA (see table 2). The ANOVA revealed that there was a significant main effect of cue condition, and a significant main effect of drug condition, but no significant main effect of order (table 2). Additionally various interactions occurred.

**Table 2.** Mixed model ANOVA details. *Factor* indicates the parameter of interest, *df* shows the degrees of freedom, *F* and *p* give the F- and p-values respectively. * symbol denotes interaction between factors.

| Factor | df | F | p |
| --- | --- | --- | --- |
| Intercept<br>(subject effects) | (1,108) | 499.4 | <0.001 |
| cue | (2,108) | 245.4 | <0.001 |
| drug (y/n) | (1,108) | 15.1 | <0.001 |
| order | (1,108) | 1.2 | 0.273 |
| cue*drug | (2,108) | 0.4 | 0.674 |
| cue*order | (2,108) | 3.7 | 0.027 |
| drug*order | (1,108) | 7.3 | 0.008 |
| order*cue*drug | (2,108) | 0.3 | 0.713 |

Alcohol intake increased attentional shift times for all 3 cuing conditions, with no interaction effect between drug and cue condition (Table 1, 2, and Figure 2). This indicates that the negative effect of alcohol (the slowing) on attentional shift times does not depend on the cuing condition (pre-, exogenous or endogenous).

The absence of a main effect of order (p=0.273, mixed model ANOVA, table 2) might indicate that shift times did not differ between the first and the second session. However, this is complicated by the finding that alcohol increased attentional shift times, and hence any learning (improvement) between session, might be concealed in the group that was under the influence in the second session. This argument is supported by the significance of the drug* order interaction (mixed model ANOVA: p=0.008, table 2). Subjects usually had faster shift times in the second compared to the first experimental session, but only if the first experimental session was the alcohol session (figure 3).

**Figure 3:**
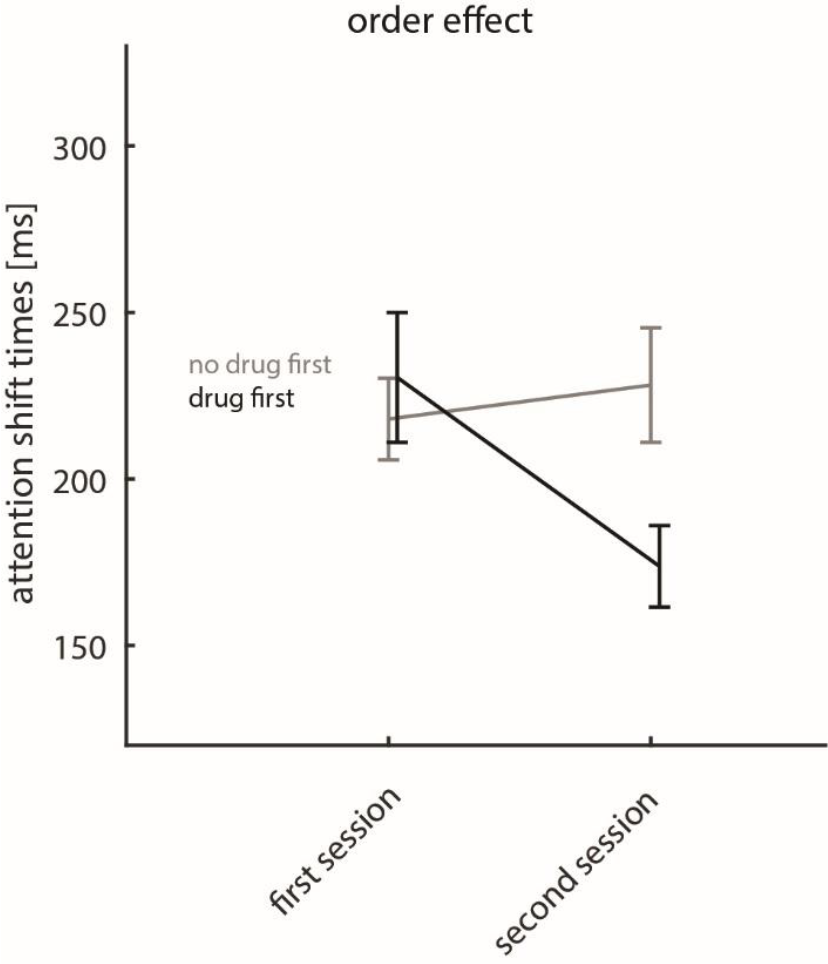
Average attentional shift times (across the 3 conditions) as a function of session (first, second) and as a function of whether they were under the influence of alcohol (drug first, black) in the first session, or whether they were not under the influence of alcohol in the first session (no drug first, grey). Session type is indicated along the x-axis, mean attentional shift times along with S.E.M are indicated along the y-axis.

Figure 3 shows that subjects’ shift times were comparatively slow when they were under the influence of alcohol in the first session, and were faster in the second session when they were not under the influence of alcohol. While group averages across attention conditions appeared marginally faster if the first session was performed under placebo conditions, this difference was not significant between groups (p=0.614, t-test, population level analysis with relatively large individual differences and relatively small sample size). However, group differences were significant for the second session, whereby subjects not under the influence of alcohol had significantly faster attention shift times than subjects that were under the influence (p=0.017, t-test). We also found a significant cue* order interaction (mixed model ANOVA, p=0.027, table 2). Changes in shift times with learning occurred for all 3 attention conditions, but they were overall largest for the pre-cue condition (Figure 4).

**Figure 4:**
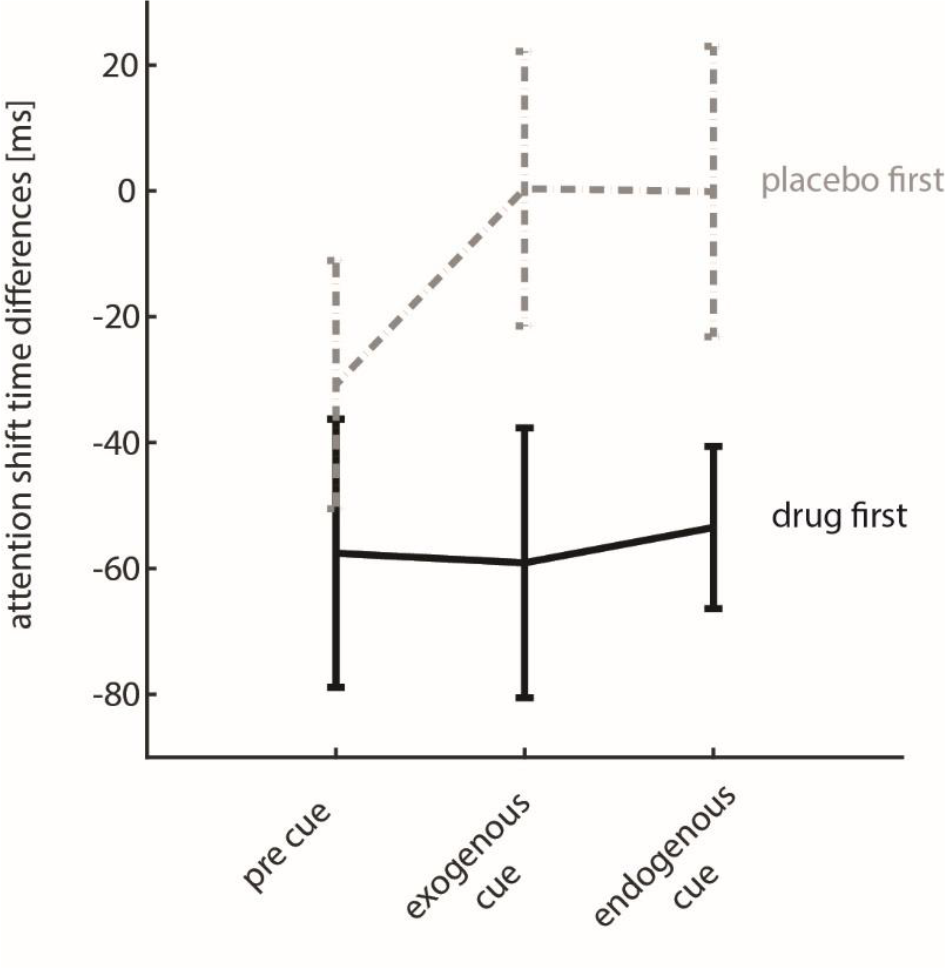
Average change in attentional shift times between session (shift times first session minus shift times second session) when placebo was taken first (grey, dashed line) and when alcohol was taken first (black line). Cue type is indicated along the x-axis, mean attentional shift time difference along with S.E.M are indicated along the y-axis.

Figure 4 shows that shift times became faster with learning, but this differed depending on whether alcohol was taken in the first or in the second session. The absolute values depicted in Figure 4 are likely a combination of the alcohol induced slowing and the speeding up that occurs with learning in the second session. To determine a baseline effect of learning, we used a dataset of 10 additional individuals (who were tested in an unrelated investigation). These subjects performed 2 sessions under identical experimental conditions (sessions were performed on separate days, at comparable times of the days, 150 trials each session, 50 pre-cue, 50 exogeneous cues, 50 endogenous cues), but they took neither drug or placebo before the session. However, for reasons unrelated to this study, they took a placebo (two small glucose pills) after the first session. The results from these sessions are shown in figure 5. For all cueing conditions shift times were faster on day 2 compared to day 1, and the effect of ‘DAY’ was significant (repeated measures ANOVA, with factors: DAY, F(1,9)= 5.7460, p=0.040; CUETYPE, F(2,18) =115.83, p<0.001; no significant interactions). The learning induced improvement in attentional shift times were about 15 ms across conditions (details Figure 5). Based on this, in conjunction with the data shown in Figure 4, it can be estimated that the alcohol induced slowing of shift times was about 30-40ms overall.

**Figure 5:**
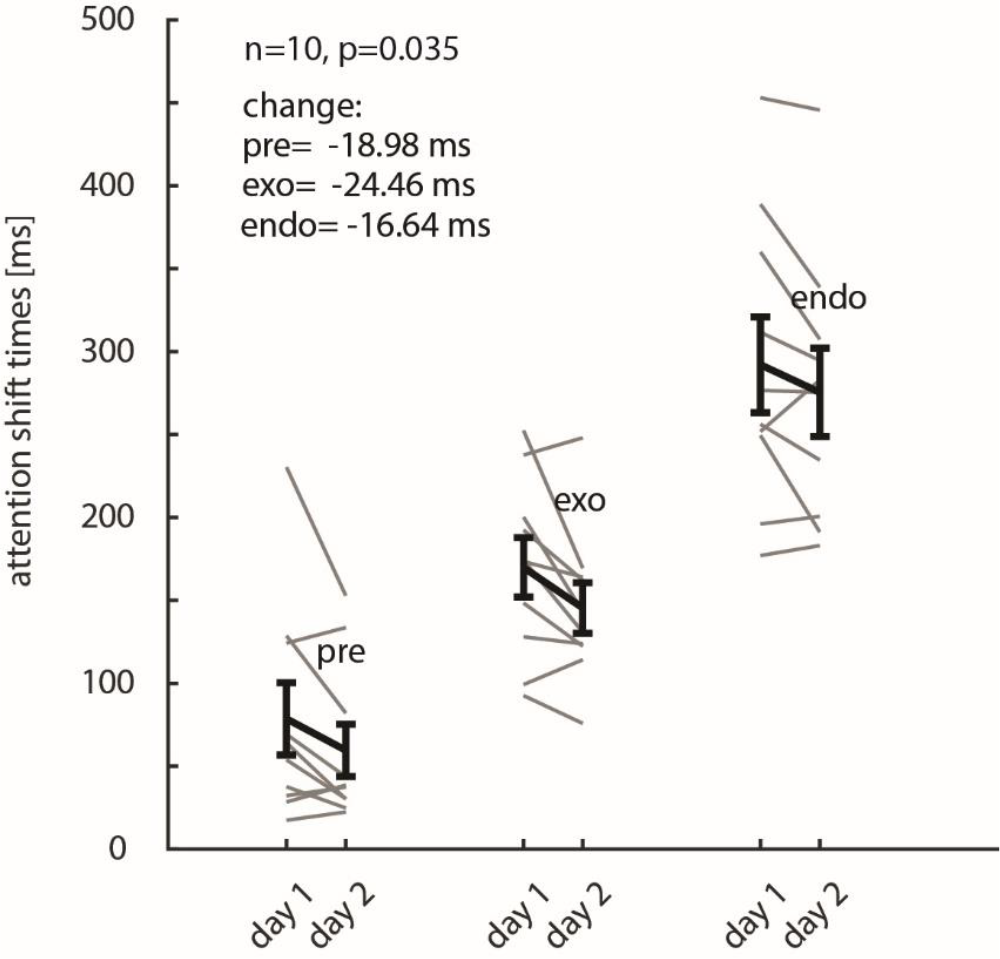
Average attentional shift times (across the 3 conditions) as a function of session (day 1, day 2) for 10 control subjects, who took neither drugs nor placebo before the sessions. Mean attentional shift times along with S.E.M are indicated along the y-axis.

### Differences in shift times as a function of Breath alcohol level

As stated previously, for each participant we measured BrAC before and after the study. The measures taken varied more than intended (range 130-360 microgram per litre breath). A simple linear regression was calculated to see if a participant’s recorded BrAC after the study predicted the difference in attentional shift latency between alcohol and placebo conditions. The relationship was not significant (*F*(1,18) =0.977, *p* = 0.336). Participants’ breathalyser results did not significantly predict how much the participants’ latencies slowed down in the alcohol condition compared with the placebo condition.

## Discussion

Our data reveal a significant increase of alcohol and on attentional shift times, compared to placebo conditions. The slowing of attentional shift times occurred for all 3 cuing conditions. Thus, alcohol induced slowing was not specific to different types of attention.

In line with previous reports (Carlson et al., 2006; Chakravarthi & VanRullen, 2011) we found that shift times were fastest for the pre-cue conditions, followed by exogenous shift, and with endogenous shift being slowest.

Our data are in line with previous studies which report negative effects of alcohol on cognition, including divided and covert attention (Schulte et al., 2001), four-choice serial reaction time, visual search tasks (Maylor & Rabbitt, 1987), number pairs and working memory (Benson et al., 2019), and inhibitory control (Fox, 1995; Klein, 2000).

Results from previous studies investigating inhibitory control (Abroms et al., 2006) or exogenously cued attention shifts (Post et al., 2000), suggested potentially differential effects of alcohol on top-down (voluntary) and bottom up (automatic) attention. Based on this study, it could have been predicted that top-down attention shift times would be increased by alcohol, while bottom-up (automatic) attention shift times would be less affected by alcohol. A similar prediction would also be made based on the global-slowing hypothesis (Maylor & Rabbitt, 1993). Our data do not support this prediction, but show a general slowing of cognition (and/or perception) irrespective of the type of attention involved.

We did not find a main effect of order on attentional shift times, which means there was no consistent change of shift times between session. However, there was an interaction between order and drug condition, as well as an interaction between order and cue condition. The absence of a main effect of order was probably due to the fact that learning which occurred between sessions (figure 5) was concealed by taking alcohol in the second session, which resulted in an overall slowing (counteracting the learning-induced improvement). Conversely, the large differences between sessions for subjects taking alcohol in the first session, are not only due to learning, as the alcohol-induced slowing would only be present in the first session. To what extent alcohol affects learning cannot be answered with the current design.

One limitation of the study is the difficulty in providing a convincing placebo. This study administered the alcohol in a cocktail containing strong citrus fruits, mint leaves and sugar in an effort to disguise the taste of the alcohol as much as possible. Unlike some other studies, we did not add a small amount of alcohol to the drink surface to disguise it even more. Aside from the taste of the cocktail, participants may also have been able to distinguish between the experimental and placebo conditions due to the effect of alcohol on their behaviour and cognition.

In conclusion, our study shows that moderate intoxication by alcohol reduces the speed of attentional shifts, irrespective of cuing conditions. This may be relevant to policy making on alcohol-related laws such as the legal drink driving limit. This study aimed for participants to be at the legal driving limit for BAC when they completed the task. This level of BAC was still found to cause a significant delay in attentional transfer speed, which could have a significant effect on drivers’ ability to detect and avoid or stop for hazards. The delay averaged 36ms, in which a driver would travel 1.13m at 70mph. While this does not seem a lot, it could be the difference between colliding with the car in front vs not.

## Acknowledgment

AT was funded by Wellcome Trust 093104 (AT), and the MRC MR/P013031/1.

